# Copper Containing Glass-Based Bone Adhesives for Orthopaedic Applications: Glass Characterization and Advanced Mechanical Evaluation

**DOI:** 10.1101/2020.11.19.390138

**Authors:** Sahar. Mokhtari, Anthony.W. Wren

**Affiliations:** Kazuo Inamori School of Engineering, Alfred University, Alfred NY, USA

**Keywords:** Bioactive Glass, Raman, Copper, Bone, Adhesive, Viscoelastic

## Abstract

This study addresses issues with currently used bone adhesives, by producing novel glass based skeletal adhesives through modification of the base glass composition to include copper (Cu) and by characterizing each glass with respect to structural changes. Bioactive glasses have found applications in fields such as orthopedics and dentistry, where they have been utilized for the restoration of bone and teeth. The present work outlines the formation of flexible organic-inorganic polyacrylic acid (PAA) – glass hybrids, commercial forms are known as glass ionomer cements (GICs). Initial stages of this research will involve characterization of the Cu-glasses, significant to evaluate the properties of the resulting adhesives. Scanning electron microscopy (SEM) of annealed Cu glasses indicates the presence of partial crystallization in the glass. The structural analysis of the glass using Raman suggests the formation of CuO nanocrystals on the surface. X-ray diffraction (XRD) pattern and X-ray photoelectron spectroscopy (XPS) further confirmed the formation of crystalline CuO phases on the surface of the annealed Cu-glass. The setting reaction was studied using Fourier transform infrared spectroscopy (ATR-FTIR). The mechanical properties of the Cu containing adhesives exhibited gel viscoelastic behavior and enhanced mechanical properties when compared to the control composition. Compression data indicated the Cu glass adhesives were efficient at energy dissipation due to the reversible interactions between CuO nano particles and PAA polymer chains.

## 1. Introduction

Bone tissue fractures are one of the most prevalent clinical diagnoses that often require invasive surgery procedures[1]. Therapeutic bone adhesives are widely used in various orthopaedic and trauma surgeries to repair and remodel the operational capacity of the bone tissue[2]. Bone adhesives are synthetic, self-curing, non-metallic materials that are mainly used either to stabilize prostheses, or used as bone fillers to fix damaged bone tissue[3]. Some of their most common clinical uses include applications in total hip or total knee replacements for anchoring functions where bone adhesive is implanted together with the prostheses [4]. Other common example is their application in treating benign aggressive bone tumours by regeneration of lost bone tissue created after curettage of tumours[5]. Injectable bone adhesives are prevalently used for bone augmentation procedures and treatment of damaged vertebral body (vertebroplasty and kyphoplasty) where bone adhesive is injected into cracked or broken vertebral body that is often caused by osteoporosis[6–8]. The principal steps in vertebroplasty and kyphoplasty involves injection of a paste of the bone adhesive into a cavity of the fractured vertebral body through a cannula, either directly or by the means of an inflatable balloon[9, 10]. Fig. 1a presents a schematic overview of the bone adhesives used in the vertebral body augmentation. The ultimate goal of using bone adhesive is to obtain optimal functional state of bone tissue by stabilization, surface adhesion, and ultimately the repair of the damaged bone tissue[11, 12]. Currently, the clinically available bone adhesives are only based on commercially available materials known as bone cement[7]. Despite being the current standard of care, the major clinical shortcomings of traditional bone cement still exist which have necessitated the need to develop alternate bone adhesives.

**Figure 1.**
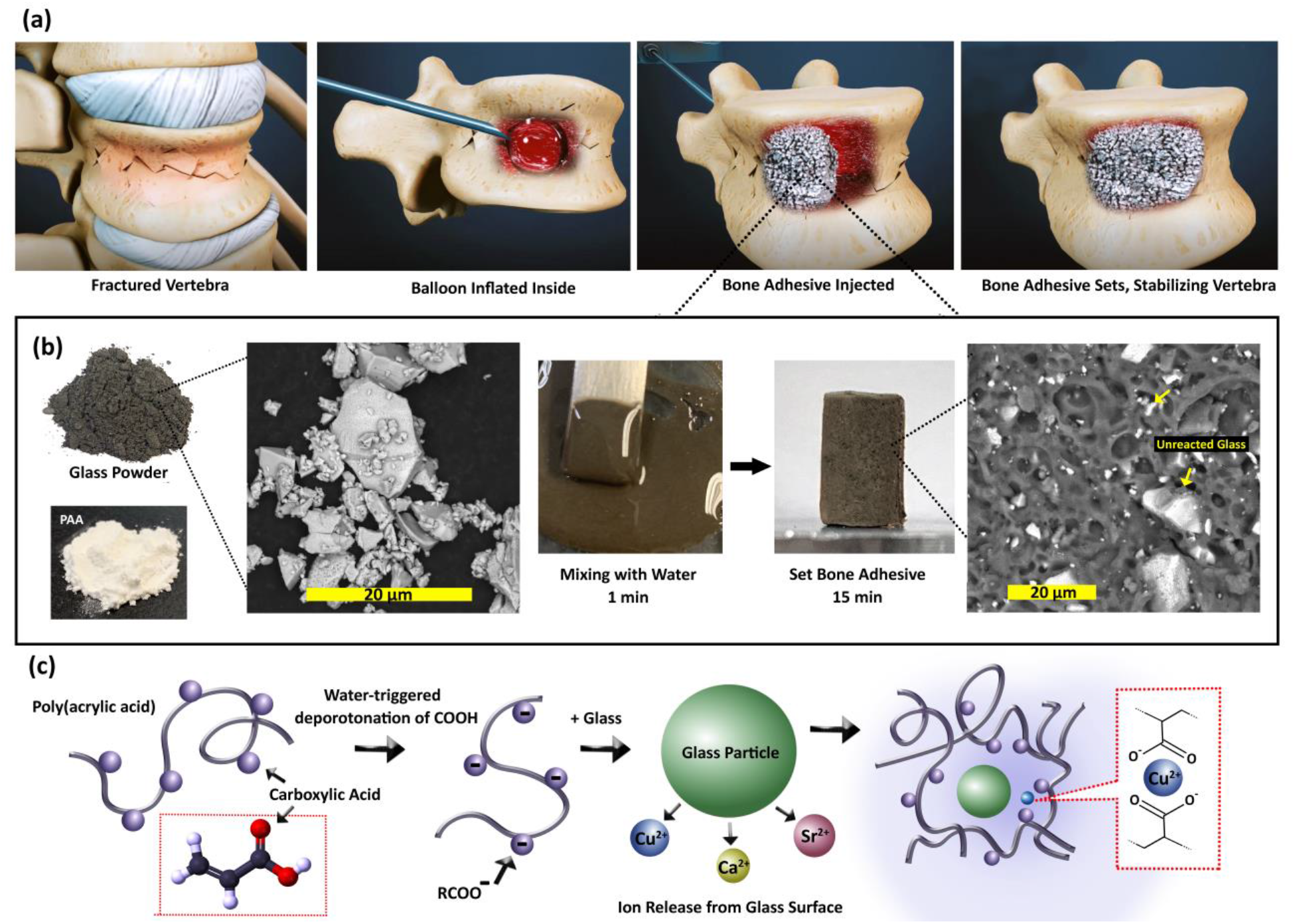
(a) Schematic illustration of the bone adhesive used in the augmentation of damaged vertebra, (b) Preparation process, where the glass powder, PAA and water were homogeneously mixed and set within 15 minutes. The SEM image shows the microstructure of the porous set adhesive. The unreacted glass particles are labeled. (c) Schematic of the setting reaction of the glass particle and polyacrylic acid in atomic level.

The first clinical bone cement used in orthopaedics, was an acrylate cement based on Polymethylmethacrylate (PMMA), which was first used for hip arthroplasty in 1958[7]. Although the clinical use and availability of various types of PMMA bone cements, they are not without major drawbacks[6, 13]. The primary concern associated with PMMA bone cement is the lack of any chemical bonding to host bone tissue which relies primarily on mechanical interlocking through penetration of cement in the irregularities in the surface of bone, acting more as cement rather than adhesive[14, 15]. Lack of interfacial chemical bonding or inadequate adhesion leads to aseptic loosening of implant, causing failure, and eventually requires revision surgeries[15]. The ideal bone adhesive should possess strong interfacial adhesion, quick and non-exothermic setting reaction, minimum swelling and volume shrinkage[1]. It is also a requirement that the bone adhesive shows biocompatibility under physiological conditions, with minimal or no cytotoxicity[16]. The unique physiochemical properties of conventionally known glass ionomer cements (GICs) could significantly benefit the field of orthopaedics; however, the mechanical properties of such materials should be addressed first[17–19]. Among different compositions of biomedical cements, the GICs showed the most biocompatibility *in-vitro*, and *in-vivo*[20, 21]. The GICs have a complex chemistry of an interpenetrating network consisting of organic and inorganic components forming a rigid crosslinked structure[19]. GICs consist of an ion-leachable glass powder (base) combined with a water-soluble polymer (acid), which upon mixing, an acid-base reaction takes place resulting in a hard set cement[22]. Since the early introduction of GICs, a number of modifications were made to these cements to improve their properties[23]. The formulation of its main constituents plays a critical role in the properties of the resultant cements. Depending on application, formulation can be modified; variations in the glass, acid, aqueous solution, and powder to liquid ratio can significantly alter setting reaction, physical, mechanical and biological behaviour[24, 25].

From mechanical perspective, a combination of strong adhesive and cohesive strength is required. Bone adhesives spread over the entire surface, evenly distribute, and transmit the physical forces throughout the contact area and effectively minimize stress localization[26]. They could also resolve certain drawbacks of metallic implants such as stress shielding, which is correlated with relatively high stiffness of metallic implants when compared to the natural bone material[15]. Natural bone tissue has a unique mechanical property and exhibits viscoelastic and rate-dependent failure behaviour. When strained, it exhibits both viscous and elastic characteristics depending on its loading condition and strain rates[27, 28]. The viscoelasticity of bone arises from its complex hierarchal structure, primarily influenced by porosity and water content[29]. Compressive experiments on femoral cortical bone over a range of strain rates, discovered that the bone would be stronger, stiffer, and more brittle at faster strain rate[29, 30].

We hereby report the fabrication of a Cu containing glass-based bone adhesive that shows exceptional viscoelastic properties. In this study, a novel glass formulation was synthesized that includes large quantities of divalent and trivalent cations in order to fully crosslink the polyacrylic acid polymer matrix, while limiting ion availability so as to allow control over the setting kinetics of the gel[31]. This property is mainly provided by inclusion of divalent ions (copper) which produce charge balanced, acid-labile tetrahedral structure, as depicted in Fig. 1. Polyacrylic acid was selected because of its biocompatibility, minimal toxicity, and excellent injectability[32]. Divalent cations in the glass composition have the capacity to link two polyanion chains, with high ionic strength of the bond. Cu^2+^ can act as a strong ionic crosslink to carboxylic acid (COO^-^) side groups, resulting in hardened set materials, which suggests the Cu is a mechanically relevant ion when formulating the adhesives[31]. In addition, Cu^2+^ has regularly been cited for being antibacterial agent[33–36]. The setting reaction in these adhesives involves three different stages: dissolution, gelation and maturation. During the first stage of the setting process, with the presence of water, the surface of glass particles is attacked by hydrogen ions from the acid chains[37]. The acid acts as proton donor and the glass powder acts as proton acceptor. Degradation starts from glass particles surface, while the core remains intact and exists as a filler in the set adhesives. Metal cations, principally Cu^2+^ and Ca^2+^ are released into solution and the pH of aqueous phase increases rapidly with release of cations[38]. This results in greater ionization of carboxylic acid and their spatial arrangement changes. Due to electrostatic repulsion, the polymer chains become more polar and uncoil, and finally take a more linear configuration. The gel structure starts to form through weak ionic crosslinking of polyacrylate chains to form a three-dimensional network[39]. The cation concentration then increases and becomes bound to the polyanionic chains. The advancement of the reaction of metallic cations with carboxylate groups leads to an increase in viscosity. This represents the initial set of the adhesives. The final material consists of unreacted glass particles surrounded by the polysalt matrix containing crosslinks. After initial hardening, further reactions take place slowly during maturation, which the less mobile cations, mainly Zn, become bound within the adhesives matrix, leading to more rigid crosslinking between the polyalkenoic chains[40–42]. The goal of this research is to address the limitations of currently used adhesives for skeletal tissue repair, and to produce novel viscoelastic glass-based adhesive that are tailored to encourage bone bonding and to improve the bioactivity of adhesives.

## 2. MATERIALS & METHODS

### 2.1 Synthesis of Glass Powders

Two glass compositions were formulated for this study, a Cu containing glass (*Cu-BG*) in addition to a Cu free control glass (*Control*). The *Cu-BG* glass contain 12 mol% CuO at the expense of Silica (SiO_2_) (Table 1). The powdered mixes of analytical grade reagents (Fisher Scientific, PA, USA) were oven dried at 100°C for 1 hour, melted at 1350°C for 3 hours in platinum crucibles, and shock quenched into water. The resulting frits were dried, ground and sieved to retrieve glass powders with a maximum particle size of 45μm. Glass powders were then annealed at 10°C/min heating profile below their T_g_ for 3 hours and cooled down to room temperature at 5°C/min.

### 2.2 Fabrication of Glass/PAA Hybrid Adhesives

adhesives were prepared by thoroughly mixing the glass powders with polyacrylic acid (E11 PAA—Mw, 210,000, <90 μm, Advanced Healthcare Limited, Kent, UK) and distilled water on a glass plate. The adhesives were formulated with a powder to liquid (P:L) ratio of 2:3 with 40 wt% additions of PAA.

### 2.3 Characterization

#### Scanning Electron Microscopy & Energy Dispersive X-ray Analysis (SEM/EDS)

Imaging was carried out with SEM (Quanta 200, FEI Company, Hillsboro, OR). Additional compositional analysis was performed with an EDAX Genesis Energy-Dispersive Spectrometer (EDS).

#### X-Ray Diffraction (XRD)

Diffraction patterns were collected using a D2 PHASER (Bruker Corporation, Billerica, MA), and phase ID was conducted using Diffrac.EVA software (Bruker Corporation, Billerica, MA). Diffractograms were collected in the range 10°<2θ<80°, at a scan step size 0.02° and a step time of 10s.

#### X-Ray Photoelectron Spectroscopy (XPS)

The X-ray photoelectron spectroscopy was carried out using a PHI (Physical Electronics, Minnesota, US) Quantera Scanning X-ray Microprobe, a monochromatic Al kα radiation (hv=1486.6 eV) at an output of 25.5 watts. Cu2p high resolution scans were collected with a pass energy of 26 eV, step size of 0.05 eV, and beam dwell time of ~ 300 ms to yield a signal to noise ratio > 100:1. Analysis area for each sample is ~ 2 to 3 mm in diameter using a 100 μm beam. Spectra analysis was performed on CasaXPS (Casa Software Ltd.). Peak positions were calibrated through normalization of the C1s peak to 284.6 eV.

#### Raman Spectroscopy

Raman analysis was performed using an Alpha300 R – Confocal Raman spectrometer (Alpha300 R, WITec, Germany) in the wavenumber range between 200 cm^−1^ to 1200cm^−1^ using a continuous wave diode laser with an excitation wavelength of 488 nm and power of 1 mW. The excitation source was focused on the sample surface using a Nikon CF Plan ELWD 50x microscope objective. Data were collected using 100 accumulations, integration time of 1 s/accumulation, and an optical grating of 1800 groves/mm.

#### ATR-FTIR Spectroscopy

The infrared spectra were recorded on a Bruker Invenio R – FTIR Spectrometer with an A225/Q-Pt ATR Multiple Crystals CRY Diamond accessory. The spectral range recorded was from 4000 cm^−1^ to 200 cm^−1^ with a resolution of 2 cm^−1^ taking the average of 50 scans.

#### Rheological Evaluation

The working time (T_w_) of the adhesives were measured under standard laboratory conditions (Ambient Temp, 25°C). Each sample (where n = 3), was measured using a stopwatch on a clean glass plate with a sterile spatula. Each measurement was conducted under the same mixing conditions to ensure reproducibility. The setting times (T_s_) of the adhesive series was measured by lowering a 400 g mass attached to a Gilmore needle into a adhesive filed mold measuring 8×9×10mm internal diameter. Adhesives were stored at 37°C during setting and the T_s_ was taken as the time the needle failed to make a complete indent in the adhesive surface, (where n = 3).

#### Mechanical Properties

The compressive strengths (σc) of the adhesives (6×4ømm, where *n = 5*) were evaluated in accordance with ISO9917. Cylindrical samples were tested after 1, and 10 hours incubation in de-ionized water. Samples were stored in sterile de-ionized water in an incubator at 37°C. After the incubation time has expired the adhesives were removed and tested while wet on an Instron 4082 Universal Testing Machine (Instron Ltd., High Wycombe, Bucks, UK) using a 1 kN load cell at a crosshead speed of 0.1 mm/min.

## 3. RESULTS

The adhesives materials under investigation as part of this work are hybrid structures, resultant of *in-situ* crosslinking between a polyacrylic acid (PAA) matrix and novel compositions of copper-containing glasses (Cu-glasses). Thus, the first part of this paper will analyse the structural properties of the glasses as a result of Cu inclusion which is significant to evaluate changes observed in the properties of the formed adhesives. The initial step in the fabrication process was to formulate the glass composition, by tailoring the chemistry required for the setting reaction. Glasses were prepared *via* melt quenching technique to obtain parent glass powders and further post-processed for surface treatment. The physical and structural properties of these glasses were compared to Cu free annealed *Control* glass, and to as prepared *Cu-BG* before heat treatment. The annealing process of glasses was performed for 3 hours just below the glass transition temperature (T_g_) of each glass composition.

The morphology and the elemental composition of the *Control*, and *Cu-BG* glass powders before and after annealing were analysed using scanning electron microscopy (SEM) and energy-dispersive X-ray (EDX). The SEM micrographs in Fig. 2 present unpolished surfaces of glass particles, showing a distribution of glass particulates measuring approximately 40 μm in diameter with a much higher concentration of smaller agglomerated fine particles below 10 μm. SEM micrographs of the annealed *Control* glass (Fig. 2) show homogenous surface features with no microscopic evidence of phase separation nor crystallization. Similar observation was found for the Cu containing glass (*Cu-BG*) before the annealing process (Fig. 2) *i.e*. smooth surface with no evidence of phase separation, porosity or crystallization. However, surface images of annealed *Cu-BG* (Fig. 2) show different morphology, with inhomogeneous microstructure of fine crystals with high density and uniform size (~100 nm) embedded on the surface of the amorphous glass phase. EDX was used to determine the elemental composition of each glass composition. Regarding the *Control* glass, the base glass constituents were detected as Si, Zn, Ca, P, and Sr. Copper containing glasses were found to have each element present in the base control glass in addition to Cu. To further investigate the crystallization of annealed *Cu-BG* glass powder, and to identify the elemental composition correlated with surface nano crystals, combined SEM-EDX elemental mapping was performed on the surface of annealed *Cu-BG* sample. Fig. 3 shows SEM micrographs of annealed *Cu-BG* glass powder with combined elemental mapping, containing the signals of Si, and Cu. The SEM micrographs of glass powder shown in Fig. 3b, and 3c clearly exhibit a crystallized surface feature for the *Cu-BG* glass, with the crystals of cubic shape or rounded cubes measuring approximately 100 nm (Fig. 3c) in size. Fig. 3b shows two types of morphologies; one shows the crystalline surface of *Cu-BG* glass (magnified in Fig. 3c), and the other displays *Cu-BG* glass particulates (marked with arrow) scattered on the surface, which in contrast to crystallized surface, do not represent any crystallization and seem to be remained amorphous. Thus, to differentiate the elemental composition of the crystalline and non-crystalline phases, elemental mapping collected from the frame depicted in Fig.3b, and results are presented in Fig. 3d-f. Through elemental mapping (Fig. 3d-f), it is shown that Cu, and Si are not uniformly distributed on the glassy and crystalline surface; the crystalline phase on the surface is enriched in Cu but depleted of Si. An opposite observation was found for the amorphous glass particulates, which were enriched in Si rather than Cu. This is further confirmed by EDX elemental analysis, revealing that the crystalline surface of glass contained about 43 at.% of Cu, and 13 at.% of Si.

**Figure 2.**
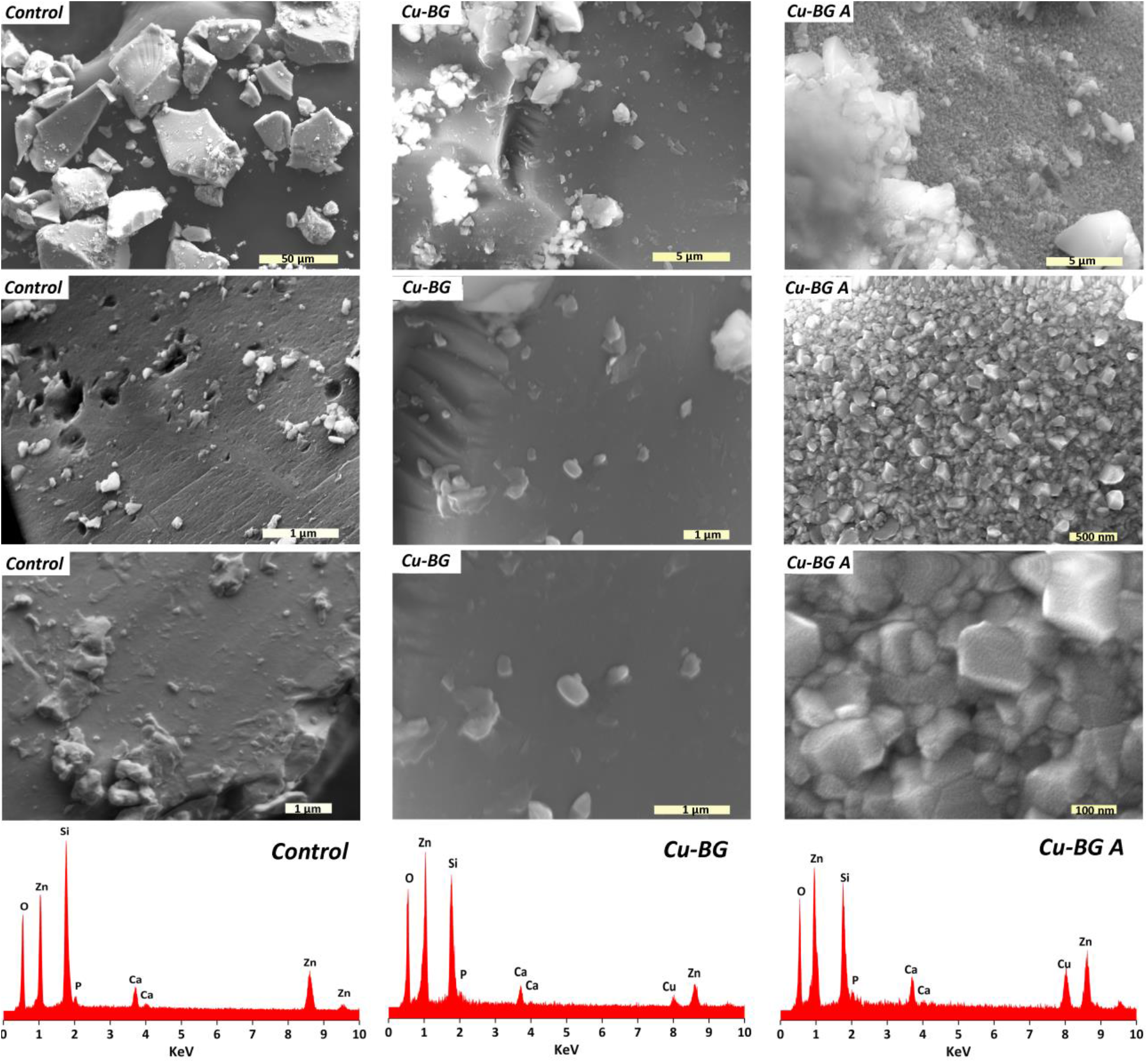
SEM images of the glass powders; *Control*, and *Cu-BG* glasses before (*Cu-BG*), and after annealing (*Cu-BG A*) with the corresponding EDX spectra.

**Figure 3.**
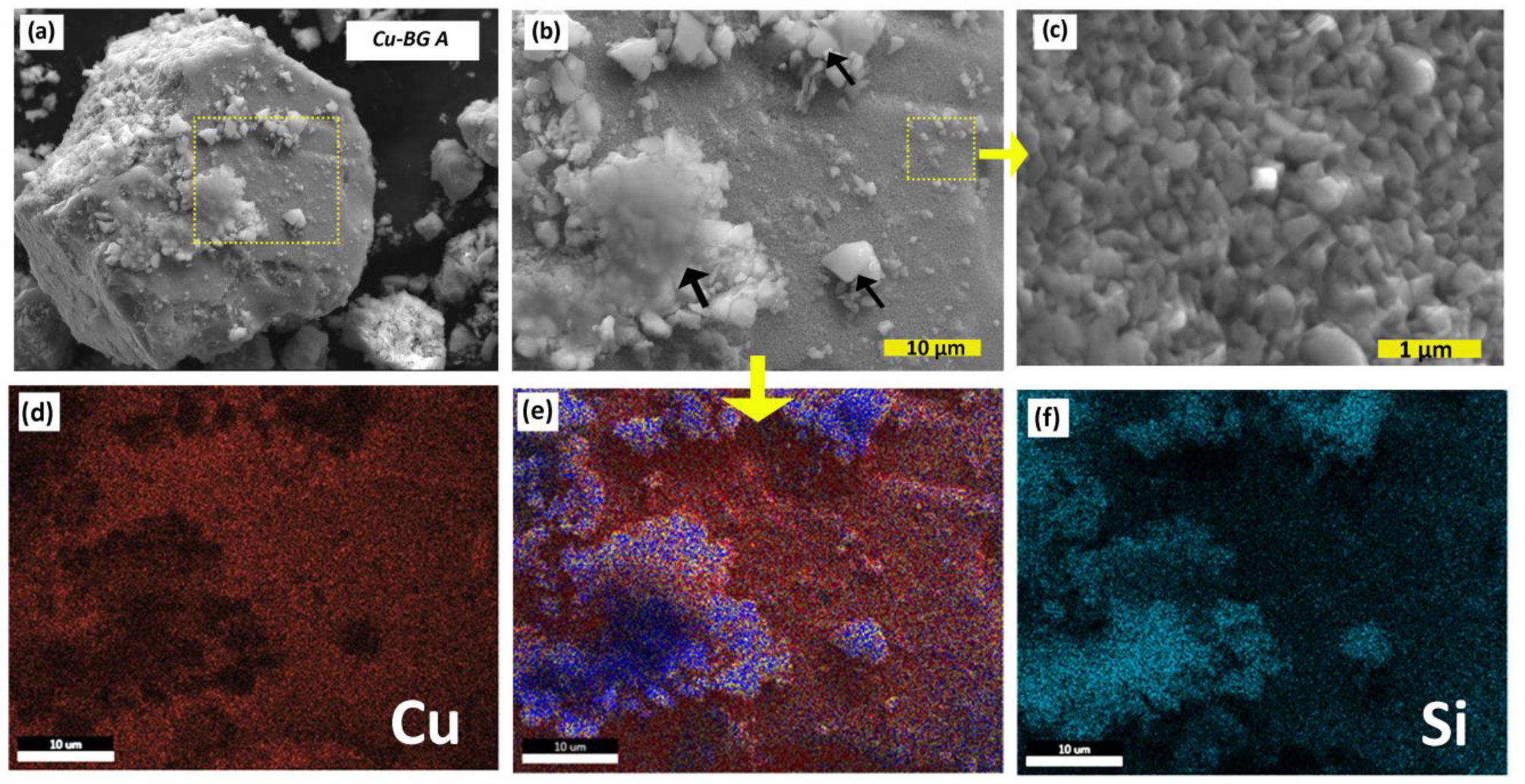
SEM images of the *Cu-BG* glass powder after annealing, and the mapping of the surface showing the distribution of Cu, and Si.

To investigate the crystalline phases observed in the SEM micrographs, glasses were examined by X-ray diffraction (XRD) analysis (Fig. 4a). A typical amorphous feature with a broad XRD halo was observed for the *Control*, and as prepared *Cu-BG* glass before annealing, confirming predominantly amorphous structures for these glasses. XRD patterns of annealed *Cu-BG* glass at 672°C, however, revealed the presence of characterise peaks corresponding to CuO crystals. An amorphous halo is still present indicating the remainder of a glassy structure for annealed *Cu-BG* powder. In order to analyse the chemical state of Cu atoms in the annealed composition of Cu containing glasses, and to what extent monovalent and divalent Cu ions exist in the glass, high resolution XPS measurements were carried out. The core-level spectra in the region of the Cu2p is shown in Fig. 4b. The two peaks at binding energies of 933 eV and 954 eV are due to the spin-orbit doublet of the Cu2p core level transition. Strong satellite peaks attributed to shake-up transition, are centred around 942 eV, and 960 eV. Each spectrum of Cu 2p3/2 and Cu 2p1/2 could be deconvoluted into two peaks. The peaks located at 932.8 eV and 953.1 eV correspond to the Cu 2p3/2 and Cu 2p1/2, respectively, indicating the presence of Cu compounds containing Cu^+^ ions in the glass. Similarly, the peaks for Cu 2p3/2 and Cu 2p1/2 located at 934.9 eV and 955.2 eV, respectively, represent the presence of Cu^2+^ ions. The results indicated that the Cu is existed in both oxidation states Cu^+^ and Cu^2+^ in the glass.

**Figure 4.**
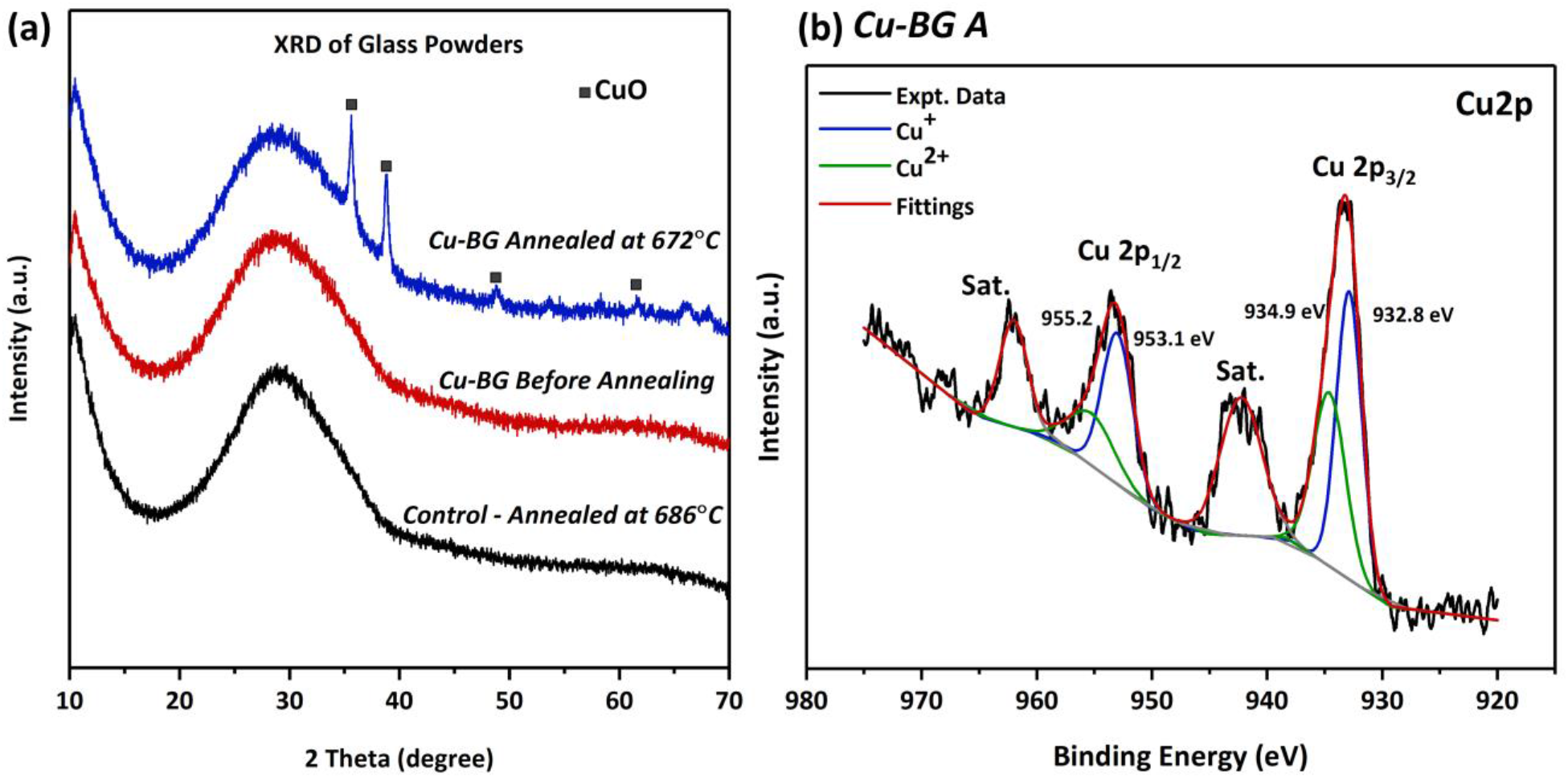
(a) XRD patterns of the glass powders, and (b) High resolution Cu2p in annealed powder of *Cu-BG* (c)

The structural characteristics of the *Control* and Cu containing glasses (*Cu-BG*) before and after heat treatment was described using Raman spectroscopy. Raman spectra of the glass powders in the frequency region between 200 and 1200 cm^−1^ are presented in Fig. 5a. All three compositions show characteristic silicon-oxygen stretching vibration observed in silicate glasses between 800-1200 cm^−1^. The band at 570 cm^−1^ is attributed to Si-O-Si bending vibration in depolymerized structural units of silicate glass. Fig. 5b shows the Raman spectrum of the crystalline phases on the surface of annealed *Cu-BG* glass powders. Three principal one-phonon modes at 282, 330, and 616 cm^−1^ in Fig. 5b, are attributed to the A_g_ and 2B_g_ modes of CuO nanostructures[43, 44]. The peak at 282 cm^−1^ is assigned to A_g_ while two other peaks at 330 and 616 cm^−1^ belong to B_g_ modes of CuO. Additionally, the widened peak at 1107 cm^−1^, is assigned to multiphonon scattering[43]. Raman spectroscopy of the annealed *Cu-BG* glass powders in Fig. 5a-b, displays two separate regions: an amorphous structure in Fig. 5a, representing high frequency bands (800-1200cm^−1^) associated with stretching vibrations of mostly depolymerised silicate species, and Raman signature of the CuO nanocrystals in Fig. 5b. The high frequency envelope (800-1200 cm^−1^) in the Raman spectra of amorphous silicate glasses, contains information about the tetrahedral silicate units and structure of the glasses[45]. To assign different type of silicate species (Q^n^) and their depolymerization degree, the vibrational spectra of glasses within the high frequency region of 800-1200 cm^−1^ were deconvoluted to four Gaussian bands attributed to vibration of the species Q^3^, Q^2^, Q^1^ and Q^0^ in the glass[46]. Results are presented in Fig. 5c-d. The band located near 850cm^−1^ is assigned to depolymerised orthosilicates (Q^0^). The region of 900–1000 cm^−1^, is associated with the Si–O stretching vibration of Q^1^ species.1100 cm^−1^, and 1000 cm^−1^ are associated with the symmetric Si–O stretching vibration of the major Q^3^, and Q^2^ species, respectively[46]. The Raman spectra of the *Control*, and *Cu-BG* glasses before and after annealing (Fig. 5c-d) indicates the existence of four different types of silicate species: Q^2^ and Q^1^ in major concentrations, and Q^3^ and Q^0^ in minor concentrations. Thus, the trend is that the more depolymerized the silica network, the lower the central frequency of the main band. In Fig. 5c-d, it can be seen that the centre of band shifts after the substitution of Cu for Si from being centred around 961 cm^−1^ for the *Control* to approximately 955 cm^−1^ for the Cu containing glasses. By deconvoluting the spectra, it can be examined to see that the substitution of the Cu leads to destruction of more polymerized units in the structure. First, the shoulder marked at 1009 cm^−1^ (Q^2^ species) loses intensity with Cu substitution alongside the growth of a new peak from what was originally a small shoulder at 860 cm^−1^ (Q^0^ species). Secondly, the shift of the most prominent peak from 1009 cm^−1^ to 943 cm^−1^ indicates the general population of silica tetrahedra is overall less connected.

**Figure 5.**
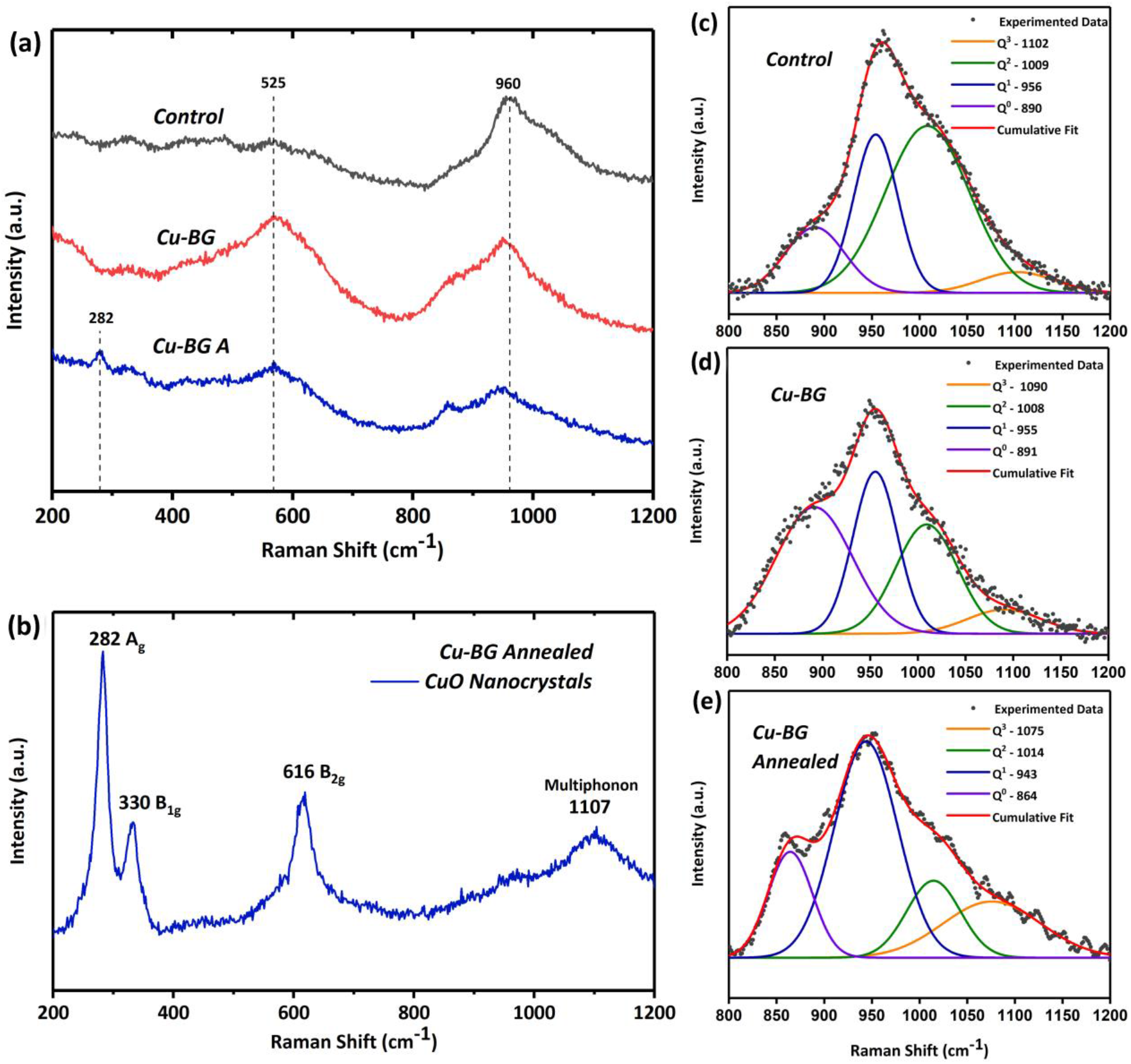
(a) Raman spectra of the glass powders and (b) Raman spectrum of crystallized *Cu-BG*, and attributed fitting curves within spectra ranges of 800-1200 cm^−1^; (a) *Control*, (b) *Cu-BG* before annealing, (c) and (d) *Cu-BG* after annealing.

Glass-based adhesives were formulated at powder to liquid ratio of 2:3, using the annealed glass powders of *Control* and *CuG* with addition of 40 wt% solution of PAA in deionized water. Hereby, the adhesives are labelled with their corresponding glass compositions as *Con/PAA* and *CuG/PAA*. Due to their similarity in the chemical composition with the glass-based ionomer cements, the working and setting time were measured according to ISO9917. The working time of the adhesives were found to be at 110s and 355s for *Con/PAA* and *CuG/PAA*, respectively. Similar trend was also observed for the setting time; 8m,46s for *Con/PAA*, and 15m,5s for *CuG/PA*. Incorporation of the Cu into the glass composition of the adhesives increased the working and setting time by 223%, and 72% respectively. To investigate the real-time molecular interactions in setting reaction of the adhesives, ATR-FTIR spectra were collected at one-minute intervals, and for over 15 minutes from beginning of the mixing. Results are presented in Fig. 6. The characteristic band for free PAA (not coordinated to a metal ion) was identified at c. 1700 cm^−1^, as was the band for ionized PAA at 1550 cm^−1^ (Fig. 6). As the setting reaction proceeded, the absorbance for the ionized PAA band increased and the absorbance for the free PAA band decreased from a clear peak into a shoulder.

**Figure 6.**
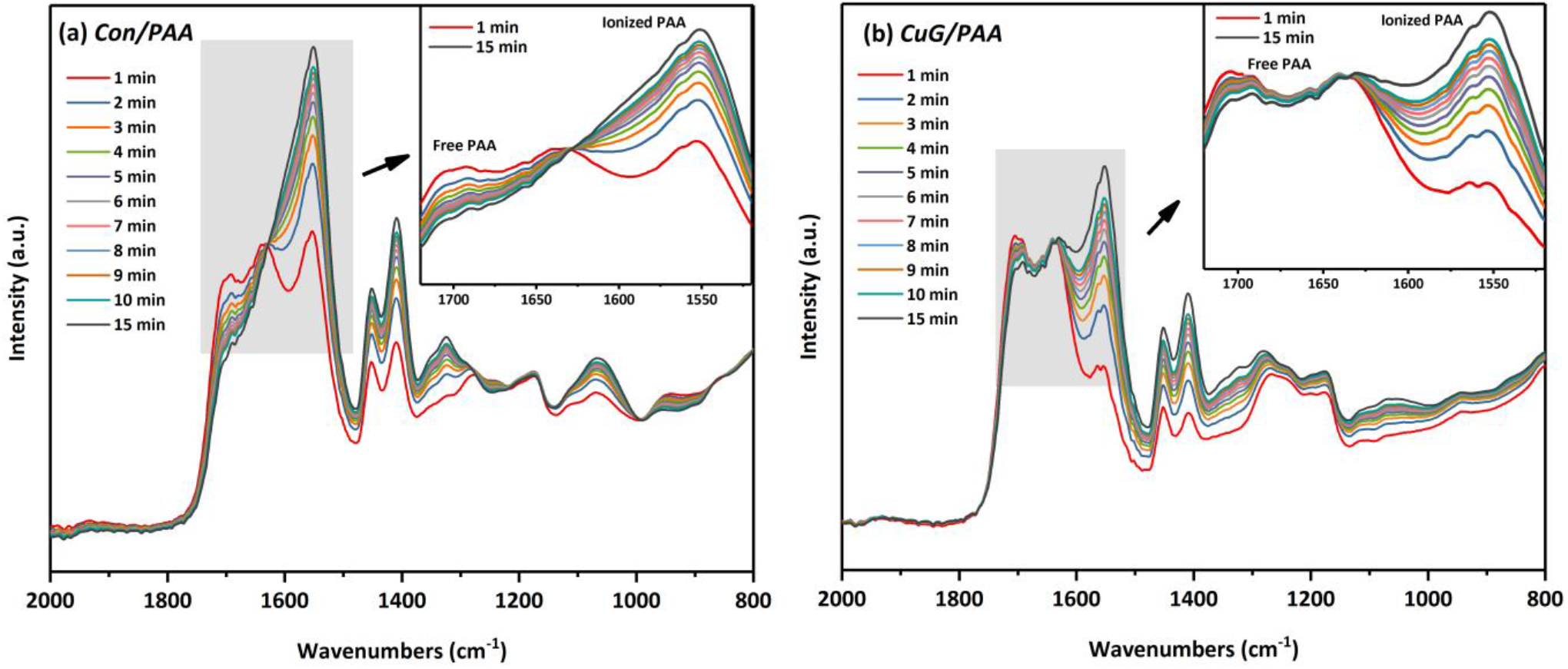
ATR-FTIR spectra within the first 15 minutes of setting reactions (a) *Con/PAA* and (b) *CuG/PAA* adhesives.

To analyse the mechanical properties of the adhesives, series of compressive measurements were performed. Fig. 7a shows a typical stress-strain curve for *Con/PAA*. The average compressive strength calculated to be at 30.7 ± 0.9 MPa. The stress-strain curve for *Con/PAA* represents the typical behaviour of brittle fracture, similar to compressive fracture of glass ionomer cements. In contrast, the compressive stress-strain curve for *CuG/PAA* in Fig. 7b, shows a different behaviour. A nonlinear viscoelastic behaviour was observed for *CuG/PAA* adhesives, with exceptional compressibility and recovery even after large compressive strains (~75%). The inset shows the sample before test, and immediately after releasing the compressive stress. Fig. 7c, and 7d show the uniaxial recovery percentage as a function of time for the *CuG/PAA* adhesives after samples were 75% compressed. The uniaxial recovery percentage was calculated as the ratio of the recovered height to the initial height of the sample. As it can be seen from Fig. 7d, the adhesives quickly recover after releasing the compressive stress. The recovery rate is approximately at 52%, 15 minutes after test, and it reaches 61% within the next 45 minutes. After this point, the recovery process slows down, and it exhibits a plateau. The recovery rate of the adhesives 48 hours after performing compressive test, was measured to be at 69% of their initial height. To better understand the viscoelastic behaviour of the *CuG/PAA* adhesives, a series of compression tests were performed to measure the compressive stress as a function of incubation time in deionized water and as a function of compressive strain (%). Results are displayed in Fig. 8, which presents the stress–strain curves of compression cycles with strain amplitude of 25, 50, and 75% for 1-hour and 10-hours incubated samples (Fig. 8a&b). The average compressive strength for corresponding strain amplitude at each time period is summarized in Fig. 8c. The compressive strength of the *CuG/PAA* adhesives after 1-hour incubation in deionized water, and at 25%, 50%, and 75% strain rates, were measured at 4 MPa, 12 MPa, and 34 MPa, respectively. For 10-hours incubated samples, the compressive strength was found to be at 6 MPa, 18 MPa, and 69 MPa, at 25%, 50%, and 75% strain rates, respectively. The stress–strain response of each loading and unloading cycle demonstrates a nonlinear rise of the stress with increasing strain, indicating nonlinear viscoelasticity of the material, and a closed loop reflecting the dissipative loss in energy in the network of the matrix[28]. The unloading curves reveals that each compression cycle results in a degree of permanent residual deformation. The stress–strain response of the adhesives changes with respect to the incubation period in water. At each value of the strain, by increasing the incubation time period of the adhesives, the compressive strength is increased. However, a higher degree of permanent deformation is also observed at each cycle for samples incubated for 10 hours. Cross section SEM micrographs of the adhesives are shown in Fig. 9. Both compositions show porous microstructure of unreacted glass particulates within the PAA matrix. The *Con/PAA* exhibits a fracture surface containing visible microcracks (Marked in Fig. 9b), and gaps at the interface of the glass particles and PAA matrix; glass particles are encapsulated within PAA matrix where there is a visible gap debonded matrix from particle creating voids. (Fig. 9c-d). The SEM micrographs of *CuG/PAA* shows a different microstructure (Fig. 9 e-h), where a homogenous matrix is observed with no visible microcracks and no interfacial disconnections between the glass particles and PAA matrix (Fig. 9h).

**Figure 7.**
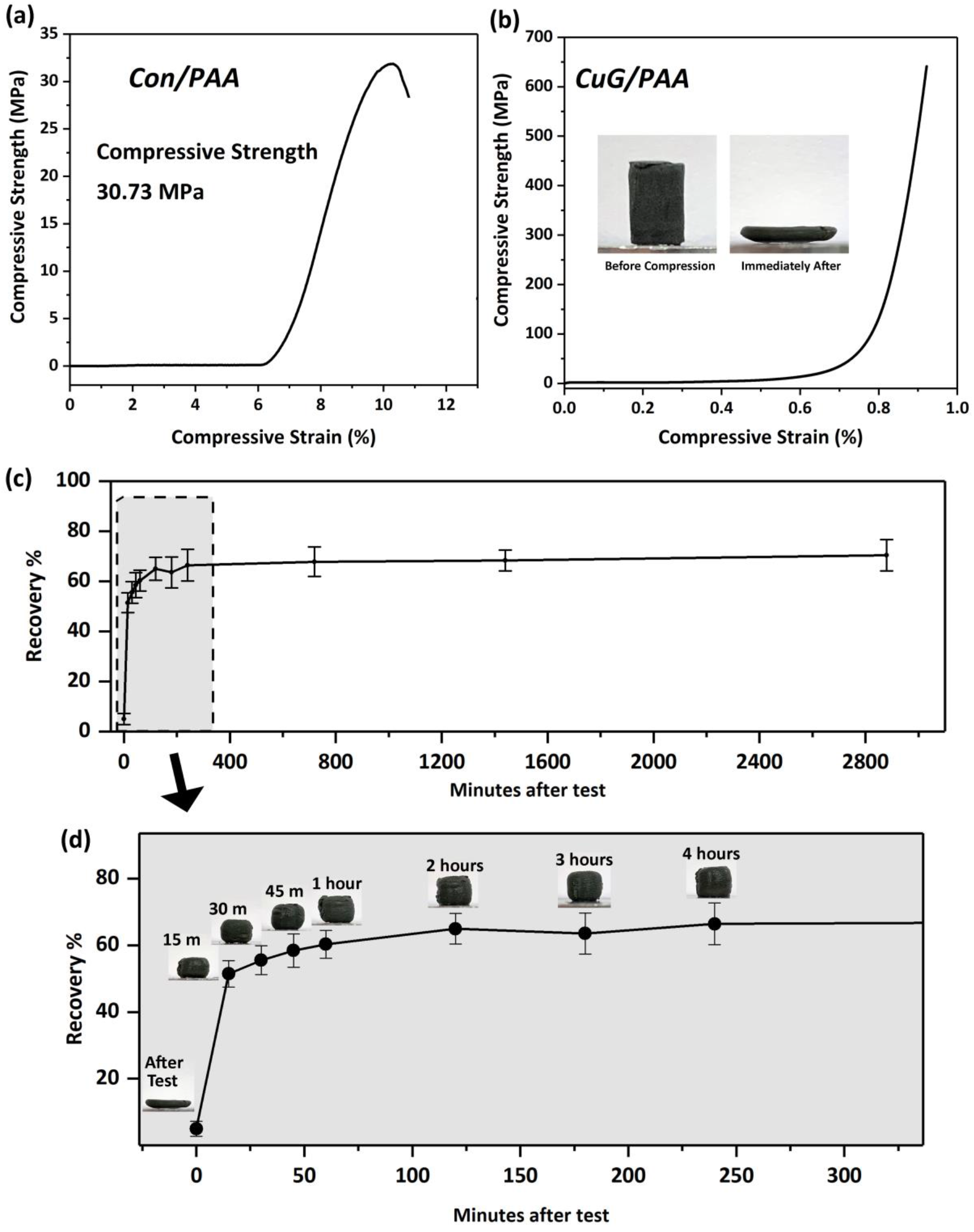
(a) Typical compressive load-extension for the *Con/PAA* and (b) *CuG/PAA* (c) Recovery of the *CuG/PAA* specimen after compression loading, (c) Projected recovery of *CuG/PAA* within the first 250 minutes.

**Figure 8.**
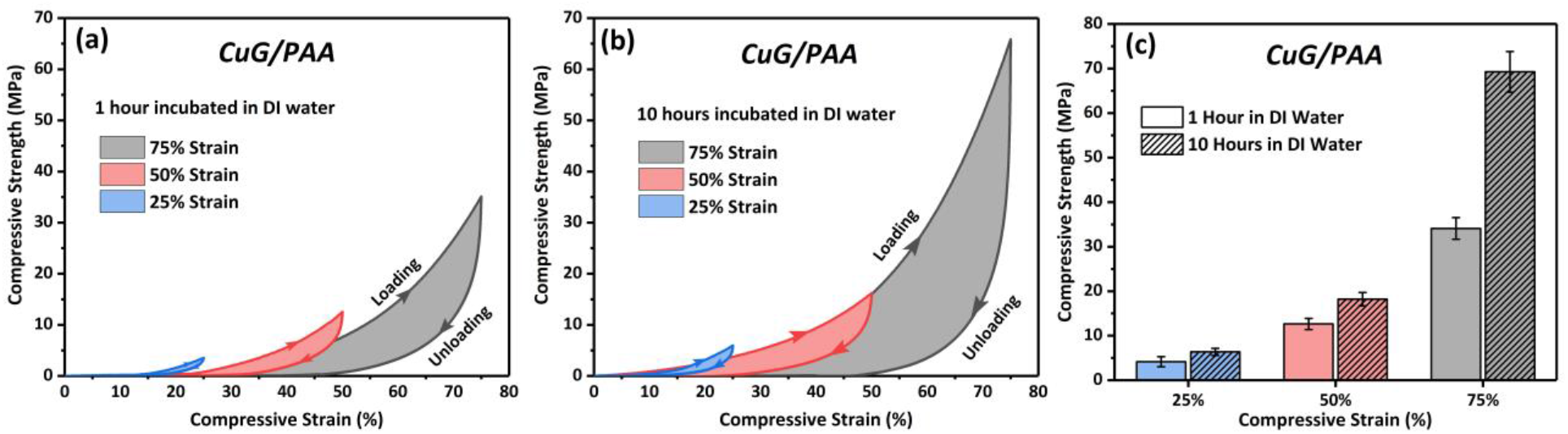
Compressive properties of *CuG/PAA* (a,b) Stress–strain curves during loading–unloading cycles in sequence of increasing strain amplitude of 25, 50, and 75% after 1 and 10 hours incubation in deionized water, (c) corresponding compressive strength of *CuG/PAA* adhesives

**Figure 9.**
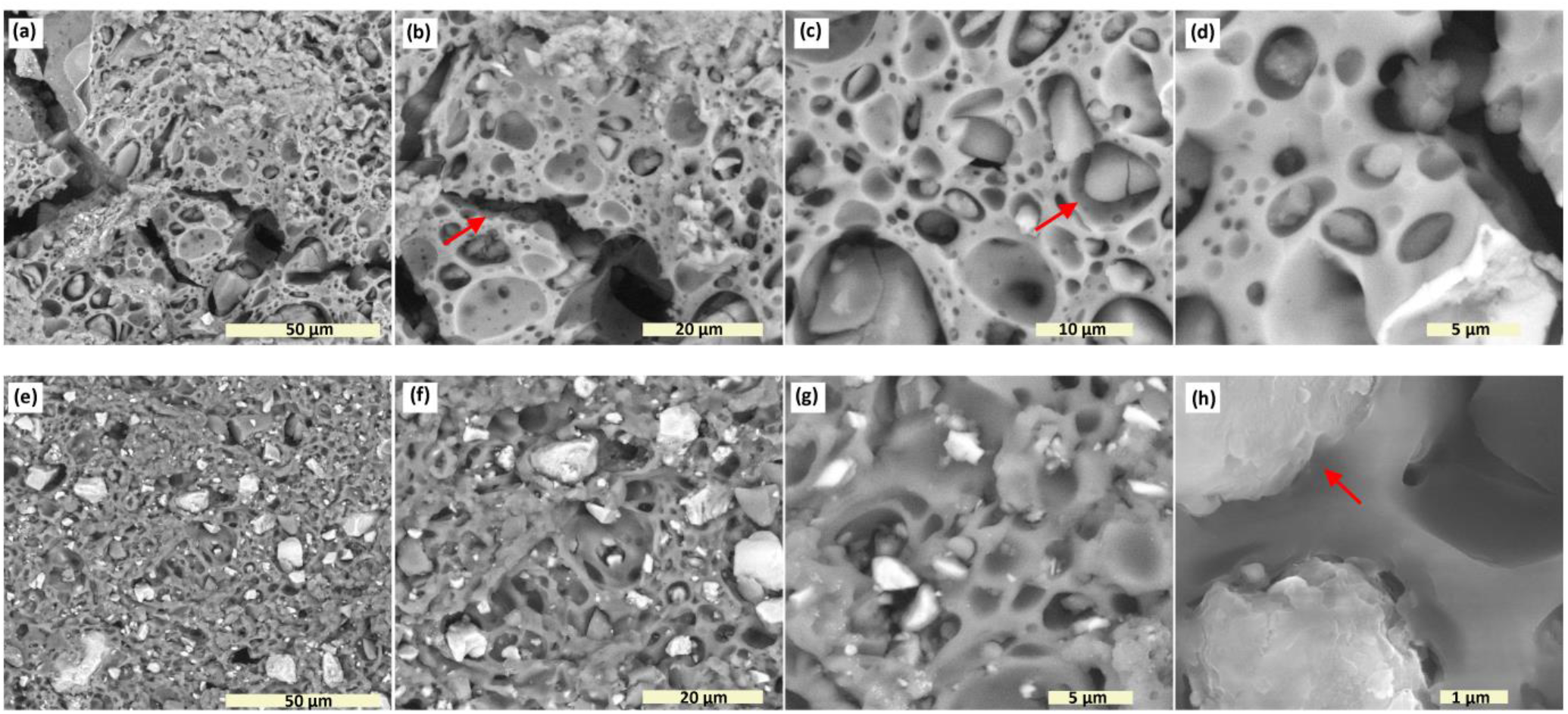
Cross section SEM micrographs of the (a-d) *Con/PAA*, and (e-h) *CuG/PAA* adhesives.

## 4. DISCUSSION

A combination of good mechanical properties, bio-adhesion, and biocompatibility is required to achieve an ideal bone adhesive[47]. The bone adhesives used for orthopaedic applications, should promote regeneration of bone tissue, encourage chemical bone-bonding and provide sufficient adhesive and cohesive mechanical strength[16]. In clinical dentistry, glass-based adhesives such as glass ionomer cements have shown exceptional properties and have been developed for various dental applications. The excellent biological performance of these materials such as bone tissue mineralization, minimal cytotoxicity, and superior biocompatibility suggests that they could provide a better and safer alternative to commercially available bone adhesives for orthopaedic applications[17]. In this study, we synthesized novel glass-based bone adhesives through modification of the glass composition and surface treatment of the glass powders. Shock quenching of the glass melt during the glass-making process introduces structural thermal stresses, which in turn influences the glass particles reactivity which affects the mechanical strength of the formed glass-based adhesive[48]. Decreased compressive strength could be attributed to the unreacted core of glass particles which acts as a reinforcing filler within the adhesive matrix[49]. When the thermally stressed glass is heat-treated to its glass transition temperature, at such viscosity, residual internal stresses in the glass structure are relieved, provided that the stress-relieved glass is cooled slowly to minimize internal stress[48]. Glass powder treated in this manner significantly improves the compressive strength of the adhesives[50]. In addition, heat treatment of the glass powders could cause a decrease in the reactivity of the glass powder through crystallization, which in turn gives the adhesives longer working and setting times and much improved handling characteristics[48]. The effects of Cu incorporation in the chemistry of these glasses, is structurally characterized in previously published studies by the authors[38, 51]. Earlier structural investigations on this glass system, suggested that the incorporation of 12 mol.% of CuO at the expense of SiO_2_ caused the glasses to undergo amorphous phase separation[38]. In this study, we reported the development of surface crystallized Cu containing glass powders, in an approach to design a glass-based bone adhesive with improved physical and mechanical properties. The SEM/EDX analysis of the glass powders presented in Fig. 2 and Fig. 3, discern surface morphologies and the elemental composition of the glass powders differentiated by composition and the heat treatment profile. The *Control* and *Cu-BG* glass powder (prior to annealing) showed no evidence of crystallization, maintaining their amorphous nature. However, upon annealing of the *Cu-BG* glass just below its Tg, the glass began to crystallize, with uniformly dispersed nano-sized cubic crystals on the surface of glass particles. Glasses show a very distinct behaviour with respect to their crystallization mechanisms[52]. The heterogeneous surface contains two phases: an amorphous glassy phase embedded with homogeneously distributed crystals[53]. Since the crystallization behaviour of the glasses is greatly influenced by the crystallization temperature in crystal growth stage, it is necessary to clarify the formation of various crystalline phases created in the glass powders at the different thermal treatment temperatures, which could influence the mechanical and structural properties of resultant adhesives.

XRD was used to identify the crystal system of nanocrystals formed upon heat treatment. XRD patterns of *control glass* (Fig. 4a) shows the presence of characteristic broad hump, confirming a predominantly amorphous glass structure. XRD pattern of the annealed *Cu-BG* glass shows an amorphous halo, and diffraction peaks corresponding to CuO, whereas it is not evident in the spectra of the as prepared *Cu-BG* glass. XRD results of *Cu-BG* glasses demonstrating partial crystallization has occurred upon heat treatment. Copper may exist in different oxidation states within glass. XPS is commonly used to identify the chemical states of constituents and to analyze transition metal complexes with localized valence d-orbitals[54] De-convoluted XPS core spectra of Cu2p3/2 and Cu2p1/2 peaks are presented in Fig. 4b. It has been reported that the shakeup satellites peaks are indicator of CuO presence at the surface[55]. As it was also shown with XRD the presence of CuO phase in the glass, XPS analysis revealed the presence of surface Cu^2+^ ions and therefore suggested that the copper compounds were predominately present at the surface as CuO nanoparticles. XRD analyses the bulk of the material, whereas, XPS measurements primarily reveal the surface information[56]. The presence of satellite peaks in Fig. 4b is an indication of compounds having Cu^2+^ ions, as it is attributed to materials having partially filled 3d^9^ shell configuration in the ground state. For metallic Cu, or Cu2O, with a completely filled shell (3d^10^), the satellite peak is reportedly absent due to the fact that the ligand-to-metal 3d charge transfer is not allowed[55]. Thus, in our prepared nanoparticles too, we are affirmative of obtaining CuO phase, formed on the outer amorphous structure of the glass particles.

Different Raman signatures involve different silicate networks around the origin[46]. The Raman spectra focuses on the band between 800 cm^−1^ and 1200 cm^−1^ where the silica tetrahedra display distinct characteristic vibrations in accordance with their network connectivity[57]. Deconvolution of the band between 800 cm^−1^ and 1200 cm^−1^, indicated that the incorporation of the modifier cations such as Cu into the structure disrupts the Si–O connectivity, resulting in the creation of non-bridging oxygens (NBOs) within the silicate structure. The notation Q^n^, where n is the number of bridging oxygens, is frequently used to distinguish between the different tetrahedral species in the network: Q^3^ corresponds to silicate species with one NBO (Si2O_5_), Q^2^ to silicate species with two NBOs (SiO_3_) and Q^1^ to silicate species with three NBO (SiO_2_), and Q^0^ is the Orthosilicate with no bridging oxygen[57]. The effect of the annealing does not significantly impact the Cu glass structure, although there is an increase in the relative intensity and broadening to lower frequency of the band associated with the Si-O-Si bending vibrations[58]. *Control* and *Cu-BG* glasses exhibited only the bands associated with the silica tetrahedra vibration, however, the Raman spectra of the annealed *Cu-BG* (Fig. 5b) glasses show the evidence of Cu-O vibration bands, as well. We observed two different domains of Raman vibrations in annealed Cu-glasses indicating of an amorphous matrix with the formation of CuO crystals on the surface of the glass[44]. Raman spectrum of annealed Cu-glass (Fig. 5b) contains the strong intensity of vibrations of nano crystalline CuO bands if compared to those of not annealed *Cu-BG*[43]. Raman spectrum of cupric oxide (CuO) includes 12 vibration modes with only 3 modes, Ag and 2Bg, active in Raman[55, 56]. In the Raman spectrum of nano crystals (Fig. 5b), we detected the presence of all three Raman active modes located at 282, 330 and 616 cm^−1^, associated with the standard modes Ag, Bg(1) and Bg(2) respectively. This is in agreement with the previous results suggesting on formation of the Cu-rich domain in the surface and nano crystallization of CuO. The results from the structural analysis of the glasses, suggests that the Cu addition in the structure of control glass leads to formation of amorphous phase separation of Cu-rich domains, and further development of nanocrystals of CuO on the surface of the glass particles upon heat treatment.

The setting reaction of the glass-based adhesives is determined by several factors including the chemistry of the glass composition (base), the type of acid, and the powder to liquid ratio in the formulation[39]. However, the setting reaction between glass and PAA is predominantly influenced by the chemistry of the glass regarding the degradation rate, and release of multivalent cations to crosslink with polymeric acid[59]. The setting reaction of the adhesives is an *in-situ* reaction between the glass powder and the PAA solution which takes place immediately upon mixing different components[60]. Hence, it is very important to characterize the working and setting times of these adhesive as these properties are significant to their clinical applicability. The setting profile was plotted as the ratio of the absorbance for the ionized PAA to free PAA over time, as described in Fig. 6. The gel formation seemed to occur within several minutes, judging from the setting time of the adhesive[61]. For both adhesive compositions the conversion ratio followed a smooth increase over time, with the rate of conversion increasing in *Con/PAA*. This trend was also observed for the final conversion ratio at 15 min. The Cu containing adhesive *CuG/PAA* (Fig. 6b) followed a different setting profile than *Con/PAA* (Fig. 6a), with a smooth increase in the conversion ratio (Fig. 6b). The ionization ratio for the *Con/PAA* was 32% higher than that of the adhesive containing Cu. These FTIR results are consistent with noted increases in working and setting times with increasing Cu content. The setting mechanism in glass-based adhesives is an acid-base reaction[62]. The reaction involves three different stages: dissolution, gelation and maturation[19]. During the first stage of the setting process, with the presence of water, the surface of glass particles is attacked by hydrogen ions from the acid chains, where the acid acts as proton donor and the glass powder acts as proton acceptor[23]. Degradation starts from glass particles surface, while the core remains intact and exists as a filler in the set cement[23]. In case of annealed *Cu-BG* glasses, the degradation is slowed down due to surface crystallization and therefore the setting reaction between glass particle and the acidic solution is delayed.

The mechanical properties of the glass-based adhesives are well understood by virtue of the formation of a continuous crosslink in the network. It is considered that the interactions between polymer-polymer, polymer-filler, and filler-filler contribute to the mechanical properties of adhesives. The high compressive strength is related to the high cross-linking density. The rigid network is mainly ascribed to the strong covalent interactions of COO^-^ chains and metallic cations released from glass particles[63]. The control adhesive of *Con/PAA* exhibits a typical brittle fracture (Fig. 7a) at low stress of the stress–strain curve[64]. This yielding phenomenon reveals the network structural changes beyond a certain strain, which is related to plastic deformation[64]. Owing to chemically covalently cross-linked networks in *Con/PAA*, a rigid microstructure with a clear yielding point was observed. Also, in the SEM micrograph of the fractured surfaces of the *Con/PAA* (Fig. 9a-d), there are more glass particles–matrix debonded sites visible due to the weakness of the glass–matrix interface compared to *CuG/PAA*. The glass particle–matrix interface is considered to be the weakest component of glass-based adhesives which may act as a stress concentration center and decreases mechanical properties where, the fracture is primarily occurred in the glass particle–matrix interface[64, 65]. However, the *CuG/PAA* shows a different mechanical response under compressive strains exhibiting unique nonlinear viscoelastic behaviour. By homogeneous distribution of CuO nanocrystals on the surface of the glass, multifunctional cross-links occurs at the interface with PAA matrix due to the combination of Cu^2+^/COO^-^ covalent cross-links (related to the plastic properties) and physical CuO/COO interactions (related to the viscoelastic properties)[66]. The debonding and rebonding of the physical cross-links such as hydrogen interactions are an effective means to dissipate deformation energy[66, 67]. The high flexibility of the *CuG/PAA* adhesives, arises from its unique crystalline surface with the reversible rearrangement of Cu/COO, resulting of efficient energy dissipation: when a crack occurs at deformations, conformations dissipate the energy and increase crack propagation resistance[50]. The loading-unloading hysteresis loops of the adhesives shown in Fig. 8, also indicates the influence of incubation period of the adhesives. A bigger hysteresis loop means a larger energy dissipated during the compression experiments and the adhesive possesses a lower elasticity with poorer repeatability[68]. Nonlinear mechanical response of networks can change dramatically when these networks are repeatedly strained, provided the bonds are not permanently cross-linked[68]. From the loading-unloading curves (Fig. 8), we can find each compression leads to a degree of permanent residual deformation, and the recoverability of the adhesive slightly decreases as a function of incubation time. This might be due to maturation process of the adhesives where it increases the ratio of chemical crosslinking between the glass particles and PAA matrix, therefore decreasing the viscoelasticity of the adhesives. *CuG/PAA* has a pronounced tendency for self-association on recovery under high strain rates, which is advantageous for the formation of load-bearing networks within the host polymer matrix[69].

## 5. Conclusion

In summary, this work reports the viscoelastic behaviour of glass-based adhesives through surface crystallization of the glass particles. In order to establish the required properties that could offer a suitable material for glass-based bone adhesives, surface crystallization was obtained upon heat treatment in *Cu-BG*. A viscoelastic behaviour was accomplished in *CuG/PAA* adhesives during compressive measurement compared with *Con/PAA*. The *CuG/PAA* exhibits exceptional viscoelastic behaviour, which is believed that could benefit a wide range of requirements in the development of bone tissue adhesives.

## 6. Acknowledgement

Part of this material (Raman data) is based upon work supported by the National Science Foundation under Grant No. DMR-1626164.

## References

[1] Böker KO, Richter K, Jäckle K, et al (2019) Current State of Bone Adhesives—Necessities and Hurdles. Materials 12:3975. doi: https://dx.doi.org/10.3390_2Fma12233975

[2] Augat P, von Rüden C (2018) Evolution of fracture treatment with bone plates. Injury 49:S2–S7. doi: https://doi.org/10.1016/S0020-1383(18)30294-8

[3] Fedak PWM, Kasatkin A (2011) Enhancing Sternal Closure Using Kryptonite Bone Adhesive: Technical Report. Surg. Innov. 18:NP8–NP11. doi: https://doi.org/10.1177_2F1553350611412057

[4] Bai S, Zhang X, Lv X, et al (2019) Bioinspired Mineral–Organic Bone Adhesives for Stable Fracture Fixation and Accelerated Bone Regeneration. Adv. Funct. Mater. 1908381. doi: https://doi.org/10.1002/adfm.201908381

[5] Marsell R, Einhorn TA (2011) The biology of fracture healing. Injury 42:551–555. doi: https://dx.doi.org/10.1016_2Fj.injury.2011.03.031

[6] Arora M, Chan EK, Gupta S, Diwan AD (2013) Polymethylmethacrylate bone cements and additives: A review of the literature. World J. Orthop. 4:67. doi: https://doi.org/10.5312/wjo.v4.i2.67

[7] Magnan B, Bondi M, Maluta T, Samaila E, Schirru L, Dall’Oca C (2013) Acrylic bone cement: current concept review. Musculoskelet Surg. 97:93–100. doi: https://doi.org/10.1007/s12306-013-0293-9

[8] Culliford D, Maskell J, Kiran A, Judge A, Javaid M, Cooper C, Arden N (2012) The lifetime risk of total hip and knee arthroplasty: results from the UK general practice research database. Osteoarthr. Cartil. 20:519–524. doi: https://doi.org/10.1016/j.joca.2012.02.636

[9] Wang W, Yeung KW (2017) Bone grafts and biomaterials substitutes for bone defect repair: A review. Bioact. Mater. 2:224–247. doi: https://doi.org/10.1016/j.bioactmat.2017.05.007

[10] He Z, Zhai Q, Hu M, Cao C, Wang J, Yang H, Li B (2015) Bone cements for percutaneous vertebroplasty and balloon kyphoplasty: current status and future developments. J. Orthop. Translat. 3:1–11. doi: https://doi.org/10.1016/j.jot.2014.11.002

[11] Kremers HM, Larson DR, Crowson CS, et al (2015) Prevalence of total hip and knee replacement in the United States. J. Bone Joint Surg. Am. 97:1386. doi: https://dx.doi.org/10.2106_2FJBJS.N.01141

[12] Mokhtari S, Wren A (2018) Bioactive glasses 2: Composite bone void fillers, Bioactive Glasses, Elsevier 2018, pp. 365–380. doi: https://doi.org/10.1016/B978-0-08-100936-9.00018-6

[13] Kim SB, Kim YJ, Yoon TL, et al (2004) The characteristics of a hydroxyapatite–chitosan–PMMA bone cement. Biomaterials 25:5715–5723. doi: https://doi.org/10.1016/j.biomaterials.2004.01.022

[14] Zhang Q-H, Cossey A, Tong J (2016) Stress shielding in bone of a bone-cement interface. MED ENG PHYS 38:423–426. doi: https://doi.org/10.1016/j.medengphy.2016.01.009

[15] Bousnane T, Benbarek S, Sahli A, Serier B, Bouiadjra BAB (2018) Damage of the bone-cement interface in finite element analyses of cemented orthopaedic implants. Period. Polytech. Mech. Eng. 62:173–178. doi: https://doi.org/10.3311/PPme.11851

[16] Farrar DF (2012) Bone adhesives for trauma surgery: A review of challenges and developments. INT. J. ADHES. ADHES. 33:89–97. doi: https://doi.org/10.1016/j.ijadhadh.2011.11.009

[17] Sidhu SK, Nicholson JW (2016) A review of glass-ionomer cements for clinical dentistry. J. Funct. Biomater. 7:16. doi: https://doi.org/10.3390/jfb7030016

[18] Akinmade A, Nicholson J (1993) Glass-ionomer cements as adhesives. J. Mater. Sci.: Mater. 4:95–101. doi: https://doi.org/10.1007/BF00120376

[19] Nicholson JW (1998) Chemistry of glass-ionomer cements: a review. Biomaterials 19:485–494. doi: https://doi.org/10.1016/S0142-9612(97)00128-2

[20] Hatton P, Hurrell-Gillingham K, Brook I (2006) Biocompatibility of glass-ionomer bone cements. J. Dent. 34:598–601. doi: https://doi.org/10.1016/j.jdent.2004.10.027

[21] Hatton P, Kearns V, Brook I (2008) Bone–cement fixation: glass–ionomer cements, Joint replacement technology, Elsevier 2008, pp. 252–263. doi: https://doi.org/10.1533/9781845694807.2.252

[22] Tyas M, Burrow M (2004) Adhesive restorative materials: A review. Aust. Dent. J. 49:112–121. doi: https://doi.org/10.1111/j.1834-7819.2004.tb00059.x

[23] Kovarik RE, Haubenreich JE, Gore D (2005) Glass ionomer cements: a review of composition, chemistry, and biocompatibility as a dental and medical implant material. J Long Term Eff Med Implants 15. doi: 10.1615/JLongTermEffMedImplants.v15.i6.80

[24] Nicholson JW, Sidhu SK, Czarnecka B (2020) Enhancing the Mechanical Properties of Glass-Ionomer Dental Cements: A Review. Materials 13:2510. doi: https://dx.doi.org/10.3390_2Fma13112510

[25] Bezerra IM, Brito ACM, de Sousa SA, Santiago BM, Cavalcanti YW, de Almeida LdFD (2020) Glass ionomer cements compared with composite resin in restoration of noncarious cervical lesions: A systematic review and meta-analysis. Heliyon 6:e03969. doi: https://dx.doi.org/10.1016_2Fj.heliyon.2020.e03969

[26] Mousa WF, Kobayashi M, Shinzato S, Kamimura M, Neo M, Yoshihara S, Nakamura T (2000) Biological and mechanical properties of PMMA-based bioactive bone cements. Biomaterials 21:2137–2146. doi: https://doi.org/10.1016/S0142-9612(00)00097-1

[27] Iyo T, Maki Y, Sasaki N, Nakata M (2004) Anisotropic viscoelastic properties of cortical bone. J. Biomech 37:1433–1437. doi:

[28] Garner E, Lakes R, Lee T, Swan C, Brand R (2000) Viscoelastic dissipation in compact bone: implications for stress-induced fluid flow in bone. J. Biomech. Eng. 122:166–172. doi: https://doi.org/10.1115/1.429638

[29] Lakes RS, Katz JL, Sternstein SS (1979) Viscoelastic properties of wet cortical bone—I. Torsional and biaxial studies. J. Biomech 12:657–678. doi: https://doi.org/10.1016/0021-9290(79)90016-2

[30] Johnson T, Socrate S, Boyce M (2010) A viscoelastic, viscoplastic model of cortical bone valid at low and high strain rates. Acta biomater. 6:4073–4080. doi: https://doi.org/10.1016/j.actbio.2010.04.017

[31] Wilson AD, Nicholson JW (2005) Acid-base cements: their biomedical and industrial applications, Cambridge University Press.

[32] Shen M, Li L, Sun Y, Xu J, Guo X, Prud’homme RK (2014) Rheology and adhesion of poly (acrylic acid)/laponite nanocomposite hydrogels as biocompatible adhesives. Langmuir 30:1636–1642. doi: https://doi.org/10.1021/la4045623

[33] Wang X, Cheng F, Liu J, et al (2016) Biocomposites of copper-containing mesoporous bioactive glass and nanofibrillated cellulose: Biocompatibility and angiogenic promotion in chronic wound healing application. Acta Biomater. 46:286–298. doi: https://doi.org/10.1016/j.actbio.2016.09.021

[34] Stähli C, James-Bhasin M, Hoppe A, Boccaccini AR, Nazhat SN (2015) Effect of ion release from Cu-doped 45S5 Bioglass^®^ on 3D endothelial cell morphogenesis. Acta Biomater. 19:15–22. doi: https://doi.org/10.1016/j.actbio.2015.03.009

[35] Li J, Zhai D, Lv F, et al (2016) Preparation of copper-containing bioactive glass/eggshell membrane nanocomposites for improving angiogenesis, antibacterial activity and wound healing. Acta Biomater. 36:254–266. doi: https://doi.org/10.1016/j.actbio.2016.03.011

[36] Manzl C, Enrich J, Ebner H, Dallinger R, Krumschnabel G (2004) Copper-induced formation of reactive oxygen species causes cell death and disruption of calcium homeostasis in trout hepatocytes. Toxicology 196:57–64. doi: https://doi.org/10.1016/j.tox.2003.11.001

[37] Prosser HJ, Powis DR, Wilson AD (1986) Glass-ionomer Cements of Improved Flexural Strength. J. Dent. Res 65:146–148. doi: https://doi.org/10.1177/00220345860650021101

[38] Mokhtari S, Skelly KD, Krull EA, et al (2017) Copper-containing glass polyalkenoate cements based on SiO2–ZnO–CaO–SrO–P2O5 glasses: glass characterization, physical and antibacterial properties. J. Mater. Sci. 52:8886–8903. doi: https://doi.org/10.1007/s10853-017-0945-5

[39] Mokhtari S, Krull EA, Sanders LM, et al (2019) Investigating the effect of germanium on the structure of SiO2-ZnO-CaO-SrO-P2O5 glasses and the subsequent influence on glass polyalkenoate cement formation, solubility and bioactivity. Mater. Sci. Eng. C 103:109843. doi: https://doi.org/10.1016/j.msec.2019.109843

[40] Wren A, Hansen J, Hayakawa S, Towler M (2013) Aluminium-free glass polyalkenoate cements: Ion release and in vitro antibacterial efficacy. J. Mater. Sci.: Mater. 24:1167–1178. doi: https://doi.org/10.1007/s10856-013-4880-y

[41] Du RL, Chang J, Ni SY, Zhai WY, Wang JY (2006) Characterization and in vitro bioactivity of zinc-containing bioactive glass and glass-ceramics. J. Biomater. Appl 20:341–360. doi: https://doi.org/10.1177_2F0885328206054535

[42] Lewis G, Towler MR, Boyd D, German MJ, Wren AW, Clarkin OM, Yates A (2010) Evaluation of two novel aluminum-free, zinc-based glass polyalkenoate cements as alternatives to PMMA bone cement for use in vertebroplasty and balloon kyphoplasty. J. Mater. Sci.: Mater. 21:59–66. doi: https://doi.org/10.1007/s10856-009-3845-7

[43] Goldstein H, Kim D-s, Peter YY, Bourne L, Chaminade J, Nganga L (1990) Raman study of CuO single crystals. Phys. Rev. B 41:7192. doi: https://doi.org/10.1103/physrevb.41.7192

[44] Xu J, Ji W, Shen Z, et al (1999) Raman spectra of CuO nanocrystals. J Raman Spectrosc. 30:413–415. doi: https://doi.org/10.1002/(SICI)1097-4555(199905)30:5_3C413::AID-JRS387_3E3.0.CO;2-N

[45] González P, Serra J, Liste S, Chiussi S, León B, Pérez-Amor M (2003) Raman spectroscopic study of bioactive silica based glasses. J Non. Cryst. Solids. 320:92–99. doi: https://doi.org/10.1016/S0022-3093(03)00013-9

[46] Yadav AK, Singh P (2015) A review of the structures of oxide glasses by Raman spectroscopy. RSC Adv. 5:67583–67609. doi: https://doi.org/10.1039/C5RA13043C

[47] Donkerwolcke M, Burny F, Muster D (1998) Tissues and bone adhesives—historical aspects. Biomaterials 19:1461–1466. doi: https://doi.org/10.1016/S0142-9612(98)00059-3

[48] Neve A, Piddock V, Combe E (1993) The effect of glass heat treatment on the properties of a novel polyalkenoate cement. Clin. Mater 12:113–115. doi: https://doi.org/10.1016/0267-6605(93)90059-G

[49] Yli-Urpo H, Lassila LV, Närhi T, Vallittu PK (2005) Compressive strength and surface characterization of glass ionomer cements modified by particles of bioactive glass. Dent. Mater. 21:201–209. doi: https://doi.org/10.1016/j.dental.2004.03.006

[50] Karimi AZ, Rezabeigi E, Drew RA (2019) Glass ionomer cements with enhanced mechanical and remineralizing properties containing 45S5 bioglass-ceramic particles. J. Mech. Behav. Biomed. Mater. 97:396–405. doi: https://doi.org/10.1016/j.jmbbm.2019.05.033

[51] Mokhtari S, Wren A (2019) Investigating the effect of Copper Addition on SiO2-ZnO-CaO-SrO-P2O5 Glass Polyalkenoate Cements: Physical, Mechanical and Biological Behavior. Biomed. Glas 5:13–33. doi: https://doi.org/10.1515/bglass-2019-0002

[52] Müller R, Zanotto E, Fokin V (2000) Surface crystallization of silicate glasses: nucleation sites and kinetics. J Non. Cryst. Solids. 274:208–231. doi: https://doi.org/10.1016/S0022-3093(00)00214-3

[53] Abo-Mosallam H, Park H (2012) Crystallization characteristics of La2O3-containing glasses for glass ionomer cement. Mater. Lett. 72:137–140. doi: https://doi.org/10.1016/j.matlet.2011.12.111

[54] Panzner G, Egert B, Schmidt HP (1985) The stability of CuO and Cu2O surfaces during argon sputtering studied by XPS and AES. Surf. Sci. 151:400–408. doi: https://doi.org/10.1016/0039-6028(85)90383-8

[55] Sahai A, Goswami N, Kaushik S, Tripathi S (2016) Cu/Cu2O/CuO nanoparticles: Novel synthesis by exploding wire technique and extensive characterization. Appl. Surf. Sci. 390:974–983. doi: https://doi.org/10.1016/j.apsusc.2016.09.005

[56] Wei B, Yang N, Pang F, Ge J (2018) Cu2O–CuO Hollow Nanospheres as a Heterogeneous Catalyst for Synergetic Oxidation of CO. J. Phys. Chem. C 122:19524–19531. doi: https://doi.org/10.1021/acs.jpcc.8b04690

[57] Notingher I, Boccaccini A, Jones J, Maquet V, Hench L (2002) Application of Raman microspectroscopy to the characterisation of bioactive materials. Mater. Charact. 49:255–260. doi: https://doi.org/10.1016/S1044-5803(03)00029-9

[58] Aguiar H, Solla E, Serra J, González P, León B, Malz F, Jäger C (2008) Raman and NMR study of bioactive Na2O–MgO–CaO–P2O5–SiO2 glasses. J. Non. Cryst. Solids. 354:5004–5008. doi: https://doi.org/10.1016/j.jnoncrysol.2008.07.033

[59] Moshaverinia A, Roohpour N, Chee WW, Schricker SR (2011) A review of powder modifications in conventional glass-ionomer dental cements. J. Mater. Chem. 21:1319–1328. doi: https://doi.org/10.1039/C0JM02309D

[60] Wetzel R, Eckardt O, Biehl P, Brauer D, Schacher F (2020) Effect of poly (acrylic acid) architecture on setting and mechanical properties of glass ionomer cements. Dent. Mater. 36:377–386. doi: https://doi.org/10.1016/j.dental.2020.01.001

[61] Valliant E, Dickey B, Price R, Boyd D, Filiaggi M (2016) Fourier transform infrared spectroscopy as a tool to study the setting reaction in glass-ionomer cements. Mater. Lett. 185:256–259. doi: https://doi.org/10.1016/j.matlet.2016.08.131

[62] de Oliveira BM, Agostini IE, Baesso ML, et al (2019) Influence of external energy sources on the dynamic setting process of glass-ionomer cements. Dent. Mater. 35:450–456. doi: https://doi.org/10.1016/j.dental.2019.01.003

[63] Charrier EE, Pogoda K, Wells RG, Janmey PA (2018) Control of cell morphology and differentiation by substrates with independently tunable elasticity and viscous dissipation. Nat. Commun 9:449. doi: https://doi.org/10.1038/s41467-018-02906-9

[64] Xie D, Brantley W, Culbertson B, Wang G (2000) Mechanical properties and microstructures of glass-ionomer cements. Dent. Mater. 16:129–138. doi: https://doi.org/10.1016/s0109-5641(99)00093-7

[65] Yamazaki T, Schricker SR, Brantley WA, Culbertson BM, Johnston W (2006) Viscoelastic behavior and fracture toughness of six glass-ionomer cements. J. Prosthet. Dent. 96:266–272. doi: https://doi.org/10.1016/j.prosdent.2006.08.011

[66] Yang J, Han C-R, Duan J-F, Xu F, Sun R-C (2013) Mechanical and viscoelastic properties of cellulose nanocrystals reinforced poly (ethylene glycol) nanocomposite hydrogels. ACS Appl. Mater. Interfaces. 5:3199–3207. doi: https://doi.org/10.1021/am4001997

[67] Chaudhuri O (2017) Viscoelastic hydrogels for 3D cell culture. Biomater. Sci. 5:1480–1490. doi: https://doi.org/10.1039/C7BM00261K

[68] Zhu C, Han TY-J, Duoss EB, Golobic AM, Kuntz JD, Spadaccini CM, Worsley MA (2015) Highly compressible 3D periodic graphene aerogel microlattices. Nat. Commun. 6:1–8. doi: https://doi.org/10.1038/ncomms7962

[69] Lei Z, Wang Q, Sun S, Zhu W, Wu P (2017) A bioinspired mineral hydrogel as a self-healable, mechanically adaptable ionic skin for highly sensitive pressure sensing. Adv. Mater. 29:1700321. doi: https://doi.org/10.1002/adma.201700321

